# Compromised ESCRT signalling is sufficient for resistance to the Target of Rapamycin Complex inhibitor Torin1 in fission yeast

**DOI:** 10.64898/2025.12.23.696270

**Authors:** John P. Alao, Rowshan A. Islam, Yusuke Toyoda, Shigeaki Saitoh, Charalampos Rallis

## Abstract

**Background:** Fission yeast cells defective in Golgi-endosomal sorting display high resistance to Torin1, a pan-Target of Rapamycin (TOR) inhibitor. TOR complexes regulate the ESCRT system to integrate nutrient availability with cell division. TOR activity is frequently deregulated in cancer, making it an attractive therapeutic target. Deregulated ESCRT activity has also been associated with cancer, but its role in mediating drug resistance is not fully understood. Herein, we have investigated the role of the ESCRT system in regulating sensitivity to Torin1.

**Methods:** Growth assays were used to monitor the growth of yeast cells. The effect of Torin1 on protein expression was monitored by immunoblotting. Fluorescence microscopy was used to investigate the action of Torin1 on protein localization.

**Results:** The ESCRT system mediates Torin1-induced degradation of amino acid and glucose transporters. The expression of these transporters at the plasma membrane is not abolished in ESCRT mutants. Mutants unable to effectively ubiquitylate these transporters are also resistant to Torin1. Impaired ESCRT-mediated protein degradation is associated with strong resistance to Torin1.

**Conclusions:** Mutations in genes encoding ESCRT components have been reported in cancer. We present evidence that compromised ESCRT signalling is sufficient for resistance to Torin1. Cells defective in ESCRT signalling or ubiquitin homeostasis are highly resistant to Torin1. Our studies demonstrate that defective ESCRT-mediated proteolysis can suppress sensitivity to Torin1.

## Background

The Target of Rapamycin Complexes, TORC1 and TORC2, integrate nutrient availability and environmental stress conditions with cell metabolism, growth and division. They, thus, play important roles in cancer development and drug resistance, and dual complex inhibitors are in clinical development. Crosstalk between the two complexes following ligand-receptor activation complicates the effective use of TORC inhibitors as anti-cancer agents. As with all anti-cancer therapeutics, *de novo* or acquired resistance limits the efficacy of pan-TORC inhibitors (1) (2). We previously reported that *Schizosaccharomyces pombe* (*S. pombe*) fission yeast mutants defective in Endosomal Sorting Complex Required for Transport (ESCRT) signalling display strong resistance to the pan-TORC inhibitor Torin1 (3). Herein, we have investigated the links between deregulated ESCRT signalling and Torin1 resistance in *S. pombe*.

The ESCRT pathway is an ancient, evolutionarily conserved pathway consisting of multicomponent complexes that execute various cellular functions, such as membrane modification, endosomal sorting, cytokinesis, membrane repair, nuclear pore sealing and viral budding in all organisms (4,5). Deregulated ESCRT activity has also been linked to genomic instability and resistance to Epidermal Growth Factor Receptor (EGFR) inhibitors and Oxaliplatin in cancer cells (6–8). In fission yeast, the TORC1 and TORC2 complexes regulate nutrient transporter localization and stability (9–11). Additionally, these complexes also regulate amino acid transporter expression by Isp7 a member of the 2-oxoglutarate-Fe(II)-dependent oxygenase gene family (12). Ubiquitylation of the Aat1 transporter by the HECT-type Pub1 E3 ligase is required for its localization to the Golgi-endosome. In the absence of Pub1, Aat1 is localized to the plasma membrane (PM). Arrestin-related molecular adaptors such as Any1 (Arn1) and Aly3 are required for Pub1-mediated ubiquitylation of nutrient transporters. When TORC1 is inhibited, or *pub1* is deleted, Aat1 localizes to the PM and is then degraded via the ESCRT pathway (9,10). Aly3 similarly regulates ubiquitylation of the Ght5 high-affinity hexose transporter. The inhibition of TORC2 abolishes Gad8-dependent phosphorylation of Aly3. This permits ubiquitylation of Ght5 by Pub1 and results in the transport of Ght5 to the endosomes for degradation in lysosomes (10,11). Inhibition of TORC2 also results in the activation of the downstream Gsk3 kinase, which is itself implicated in cancer development and progression. Gsk3 mutants are also highly resistant to Torin1. Phosphorylation of Pub1 by Gsk3 has been reported to regulate its activity (3,13). Interestingly, Pub1 also regulates Cdc25 stability, targeting it for ubiquitin-dependent proteasomal degradation. Cdc25 is a phosphatase that regulates cell cycle progression by counteracting the inhibitory phosphorylation of Cdc2 by Wee1(14). Cdc25 degradation is required for cytokinesis and proper cell division following mitosis (15). Wee1 has been reported to delay cell cycle progression in G2 following glucose withdrawal, and *wee1* mutants lose viability under these conditions (16). Inhibition of Cdc2 activity leads to arrest in the G1, S or G2 phases of the cell cycle in a context-dependent manner (14).

The expression of PM nutrient transporters is regulated by TORC1 and TORC2 according to nutrient availability. In the absence of nitrogen, cells rapidly divide and exit the cell cycle or undergo mating. In contrast, glucose limitation induces a transient cell cycle arrest in G2 as Ght5 expression is upregulated (11,16,17). Exposure of fission yeast cells to caffeine or rapamycin inhibits TORC1 but not TORC2, driving entry into mitosis without subsequent cell cycle arrest. In contrast, exposure to Torin1 inhibits TORC1 and TORC2, which drives cells into mitosis and cell cycle arrest in G2 (17,18). TORC1 and TORC2, together with Gsk3 and Pub1, thus regulate Cdc25 activity, integrating nutrient availability with cell cycle regulation. Furthermore, Torin1 induces Cdc25 degradation and cell cycle arrest due to the inhibition of Cdc2 activity by Wee1 (17,19). We hypothesized that Torin1 induces the global degradation of amino acid and glucose transporters, leading to cell cycle arrest due to Cdc2 inhibition. Mutations that confer resistance to Torin1 would need to rewire cell signalling pathways to reactivate or prevent the inactivation of Cdc2.

We report that Torin1 treatment results in the degradation of the Aat1 and Cat1 amino acid transporters as well as the Ght5 glucose transporter in an ESCRT-dependent manner. Furthermore, Pub1 is required for the degradation of these transporters following exposure to Torin1. In *pub1*Δ and ESCRT mutants, Aat1, Cat1 and Ght5 expression is not abolished in response to Torin1. Interestingly, suppression of global protein ubiquitylation also confers resistance to Torin1. We propose that fission yeast strains defective in ESCRT-mediated protein degradation maintain the expression of nutrient transporters at the PM following exposure to Torin1. Deregulated ESCRT activity can thus contribute to Torin1 resistance by preventing cell cycle exit.

## Methods

### Strains, media, and reagents

Strains are listed in Additional file 1. Cells were grown in yeast extract plus supplements medium (YES). For experiments in the absence of nitrogen, we used EMM2-N medium (Formedium, Hunstanton, United Kingdom). Stock solutions of caffeine (Sigma Aldrich, Gillingham, United Kingdom) (100 mM) were prepared in water stored at -20°C. Rapamycin (Tocris, Abingdon, United Kingdom) was dissolved in DMSO (500µg/mL) and stored at - 20°C. Torin1 (Tocris) was dissolved in DMSO (3.3mM) and stored at -20°C.

### Microscopy

Fluorescent microscopy of GFP-expressing strains was performed using an EVOS M5000 and Leica DMRA2 epi-fluorescent microscope attached to a monochrome Orca-ER camera from Hamamatsu Photonics.

### Immunoblotting

Antibodies directed against GFP (11814460001 Roche) were purchased from Sigma-Aldrich (Gillingham, United Kingdom). Secondary antibodies directed against mouse (ab205719) IgG were purchased from Abcam (Abcam, Cambridge, United Kingdom). Cell pellets were prepared for SDS-PAGE and treated as previously reported (Alao et al., 2025). Where deemed appropriate, images were quantified using Image J software and Student’s t-test for statistical analyses.

## Results

### Torin1 and nitrogen withdrawal induce Ght5 degradation

We previously identified genes encoding ESCRT pathway components in screens for Torin1 resistance (3,21). As part of this study, we also identified *hse1* (ESCRT 0) and *sst6* (*vps23*) (ESCRT I) deletion mutants as being resistant to the drug. We also identified *atg6* and *vps38* (required for the activation of ESCRT signalling) but not *atg36* (regulates the activation of autophagy) mutants in our screens, indicating that the ESCRT pathway confers sensitivity to Torin1 (Fig. 1A (Light green and dark green ovals, respectively) and Fig. 1B). *S. pombe* cells cannot efficiently degrade GFP, which accumulates as Ght5 is degraded (22,23). We thus investigated the effect of Torin1 on transmembrane nutrient transporters using strains expressing Ght5-GFP (11). Exposure to 5 µM Torin1 but not 10 mM caffeine for 3h induced Ght5-GFP degradation independently of Tor1. We noted, however, that Ght5-GFP degradation was impaired in the *tor1*Δ mutant compared to *wt* cells (See discussion) (Fig. 1C and 1D). Exposure to 100 ng/ mL rapamycin slightly induced Ght5-GFP degradation, as indicated by the accumulation of GFP. However, exposure to rapamycin induced Ght5-GFP accumulation. Exposure to caffeine and rapamycin induced Ght5-GFP degradation to an extent like that of Torin1. It is unclear if this effect is due to the induction of autophagy rather than Tor1 inhibition and degradation *via* ESCRT signalling. Caffeine also counteracted the effect of rapamycin on Ght5-GFP accumulation (Fig. 1E). While we identified *atg6* as the only autophagy-regulating gene in our screen for Torin1 resistance, exposure to the drug also induces autophagy as detected by GFP-Atg8 cleavage (Additional file 1A). In contrast to glucose deprivation, exposure to Torin1 thus mimics the effects of nitrogen withdrawal on Ght5-GFP (Fig. 1F). Microscopic analyses demonstrated that exposure to 5 µM Torin1 induced Ght5-GFP vacuolar accumulation after 1h, followed by degradation (1h vs 3h) (Fig. 1C and 1G). Similar results were observed only when caffeine was combined with rapamycin. Caffeine and rapamycin also exerted differential effects on Ght5 localization, which was more diffuse than the latter. This is consistent with their mechanisms of TORC1 inhibition. We also noted that in contrast to cells exposed to Torin1, cells exposed to caffeine with or without rapamycin were still undergoing division after 3h (Fig. 1G). Since TORC2 prevents Ght5-GFP ubiquitylation, we conclude that caffeine combined with rapamycin induces its degradation independently of the ESCRT system. In contrast, Torin1 induces the degradation of Ght5-GFP via the ESCRT pathways by inhibiting TORC2. Pub1 and Pub3 have been identified as Ght5 E3 ligases (10).

**Fig. 1.**
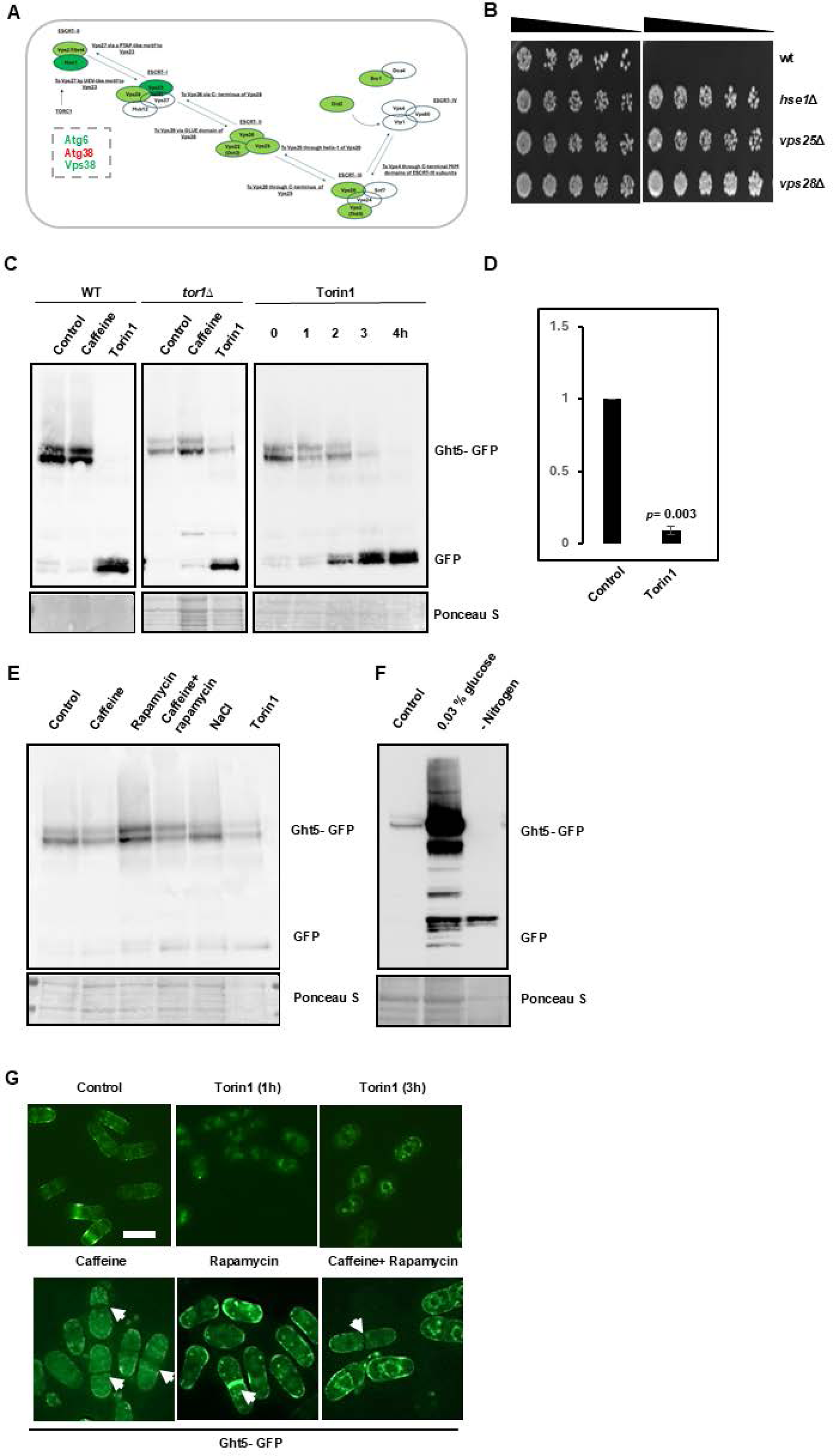
Torin1 induces Ght5 degradation. **A** Mutations in ESCRT complex protein subunits induce resistance to Torin1. Subunits in dark green were identified independently of our drug screening. The resistance of *atg6* and *vps38*, but not *atg38* mutants links ESCRT signalling to Torin1 sensitivity. **B** Wt, *hse1*Δ, *vps25*Δ and *vps28*Δ mutants were grown to stationary phase and then cultured in fresh YES media for at least 2h. Cells were adjusted to an O.D.600 of 0.3 to 0.5 and plated on YES agar with or without 5µM Torin1 and incubated at 32°C for 2-3 days. **C** Cells expressing Ght5-GFP were grown to log phase and exposed to 5µM Torin1 for 3h or as indicated. Cell pellets were stored at -80° C and analysed by SDS-PAGE. GFP expression was monitored by immunoblot analysis using monoclonal antibodies directed against GFP. Full-length Ght5-GFP and cleaved GFP is indicated. Ponceau S was used to monitor gel loading. **D** Quantification of Torin1-induced Ght5-GFP degradation. Experiments were carried out as in **C** (*n*= 3). **E** Cells were grown to log phase and incubated with 10mM caffeine, 100ng/mL Rapamycin alone or in combination, 150mM NaCl or 5µM Torin1 for 3h. **F** Cells were grown to log phase and transferred into fresh media with 3% glucose, 0.03% glucose or EMM2-N for 3h. Samples were analysed as in D. **G** Cells expressing Ght5-GFP were grown to log phase, exposed to 5µM Torin1 for 1 and 3h and examined by fluorescent microscopy. White arrowheads indicate septa. Scale bars indicate 10µm.

The deletion of *pub1* clearly suppressed the degradation of Ght5-GFP, confirming our previous findings. Ght5-GFP degradation was not abolished in this mutant, likely due to residual Pub3 activity (Additional file 1C). Torin1 thus arrests cell division in G2, in part by preventing glucose import via the Ght5 transporter at the PM (10,17).

### Defective ESCRT signaling attenuates Torin1-mediated Ght5 degradation

Next, we investigated the effect of Torin1 on Ght5-GFP expression in ESCRT mutants. In contrast to wt cells, Torin1 failed to induce Ght5-GFP degradation in *hse1*Δ mutants. We noted that this mutation did not abolish Ght5-GFP degradation as GFP still accumulated. Deletion of *sst4* did not prevent Ght5-GFP degradation, although basal levels were higher, and we did not observe the accumulation of GFP in this mutant (Fig. 2A to 2C). Exposure to Torin1 did not induce Ght5-GFP degradation in *vps25*Δ mutants. GFP still accumulated in these mutants, suggesting the degradation of Ght5-GFP is not completely abolished (Fig. 2D and 2E). Inhibition of Tor1 or Gad8 activates Gsk3, which regulates Pub1 and Pub3 activity (ref) (Fig. 2F). We deleted *gsk3* in the Ght5-GFP mutant to investigate its role in regulating Ght5 stability. However, deletion of *gsk3* did not affect Ght5-GFP expression or following exposure to Torin1 (Fig. 2G to 2I, Additional file 1B). The resistance of *gsk3* mutants to Torin1, thus, appears to be unrelated to its regulation of Pub1/ Pub3 expression and Ght5-GFP ubiquitylation/ degradation (ref, ref). Our results identify the ESCRT system as being required for Torin1-induced Ght5 degradation.

**Fig. 2.**
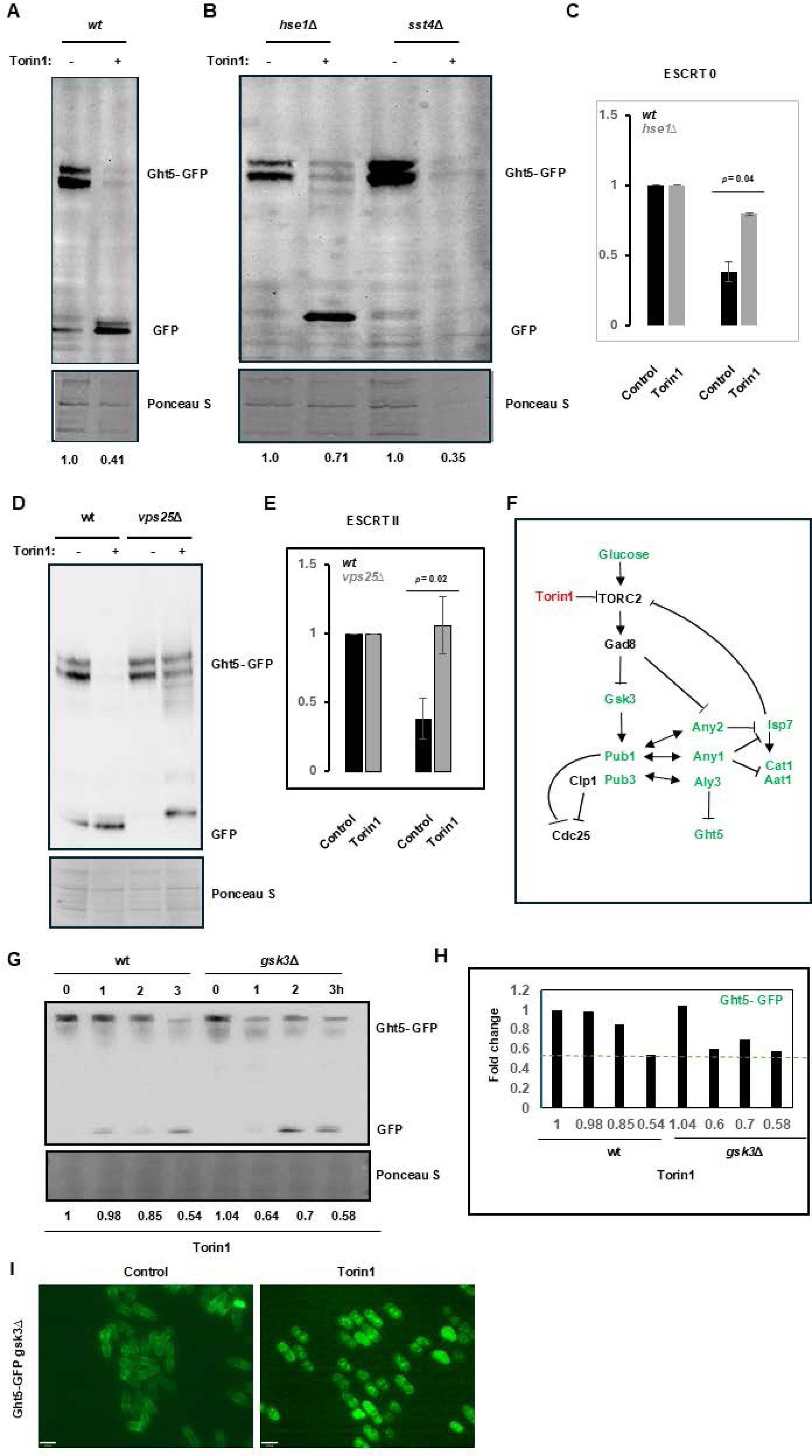
Torin1 induces ESCRT-dependent Ght5 degradation. **A** *Wt* cells expressing Ght5-GFP were exposed to 5µM Torin1 for 3h and samples analysed by immunoblot analysis. Numbers indicate relative expression levels. **B** *hse1*Δ and *sst4*Δ mutants were exposed to 5µM Torin1 for 3h and analysed as in A. **C** Quantification of Torin1-induced Ght5 degradation in wt and *hse1*Δ cells (*n*=3). **D** Wt and *vps25*Δ mutants were treated as in A. **E** Quantification of Torin1-induced Ght5 degradation in wt and *vps*Δ cells (*n*=3). **F** Regulation of nutrient transporter stability by TORC2. Torin1 inhibits TORC2, blocks Gad8 activity and induces Gsk3 activation. Gsk3 stabilizes Pub1, which targets Cdc25 for ubiquitin-dependent proteasomal degradation. Inhibition of Gad8 facilitates arrestin-related adaptor-mediated ubiquitylation of Aat1, Cat and Ght5 by Pub1 and Pub3. TORC2 also regulates Aat1 and Cat1 expression by regulating the activity of Isp7. **G** *Wt* and *gsk3*Δ mutants were exposed to 5µM Torin1 as indicated. Samples were processed as in A. **H** Quantification of Torin1-induced Ght5 degradation in *wt* and *gsk3*Δ mutants (*n*=1). **I** *Wt* and *gsk3*Δ mutants expressing Ght5-GFP were exposed to 5µM Torin1 for 3h and examined by fluorescent microscopy. Scale bars indicate 10µm.

### ESCRT mutants maintain Ght5 expression at the PM following exposure to Torin1

We next compared the expression of Ght5-GFP in wt and ESCRT pathway mutants following exposure to 5 µM Torin1. In untreated cells, Ght5-GFP was expressed at the PM in bright patches with little expression in the cytoplasm. Exposure to Torin1 induced the rapid import of Ght5-GFP in a speckled pattern and subsequent degradation within the vacuole after 3h (Fig. 1 and 3A). ESCRT mutants also expressed Ght5 at the PM, but the expression pattern differed in different mutants. Ght5-GFP was visible in a granular cytoplasmic pattern even in untreated *hse1*Δ, *vps25*Δ mutants and strongly at the cell membrane in *sst4*Δ, and *sst6*Δ cells (Fig. 3B to 3E). Defective TORC2 and ESCRT-mediated protein degradation thus result in the accumulation of Ght5 (ref). We noted by immunoblotting that basal Ght5-GFP levels were not identical in these mutants (not shown). Following exposure to Torin1, Ght5-GFP was still expressed at the PM in *hse1*Δ, *vps25*Δ, *sst4*Δ, and *sst6*Δ mutants. Torin1 exposure did not affect the expression of Ght5 in *sst4*Δ mutants (Fig. 3D). Exposure to Torin1 appeared to increase the expression of Ght5 at the PM in *sst6*Δ mutants (Fig. 3E). We concluded that, unlike wt cells, *hse1*Δ, *vps25*Δ, *sst4*Δ, and *sst6*Δ mutants continue to express Ght5-GFP at the PM following exposure to Torin1 (Fig. 3A to 3E). Glucose is required for cell division, and low concentrations strongly induce the expression of Ght5 at the PM (Fig. 1F) (11). We investigated whether growth on 0.03 % glucose induces resistance to Torin1. The Torin1 sensitivity of the *wt*, *tor1*Δ and ESCRT mutants rather increased under these conditions (Additional file 1B). Thus, overexpression of Ght5 alone is insufficient to suppress Torin1 sensitivity. Torin1 induces Ght5 degradation, and its expression is maintained at the cell membrane of ESCRT mutants. These findings partly explain the different effects of caffeine, rapamycin and Torin1 on cell cycle progression in fission yeast (17,24).

**Fig. 3.**
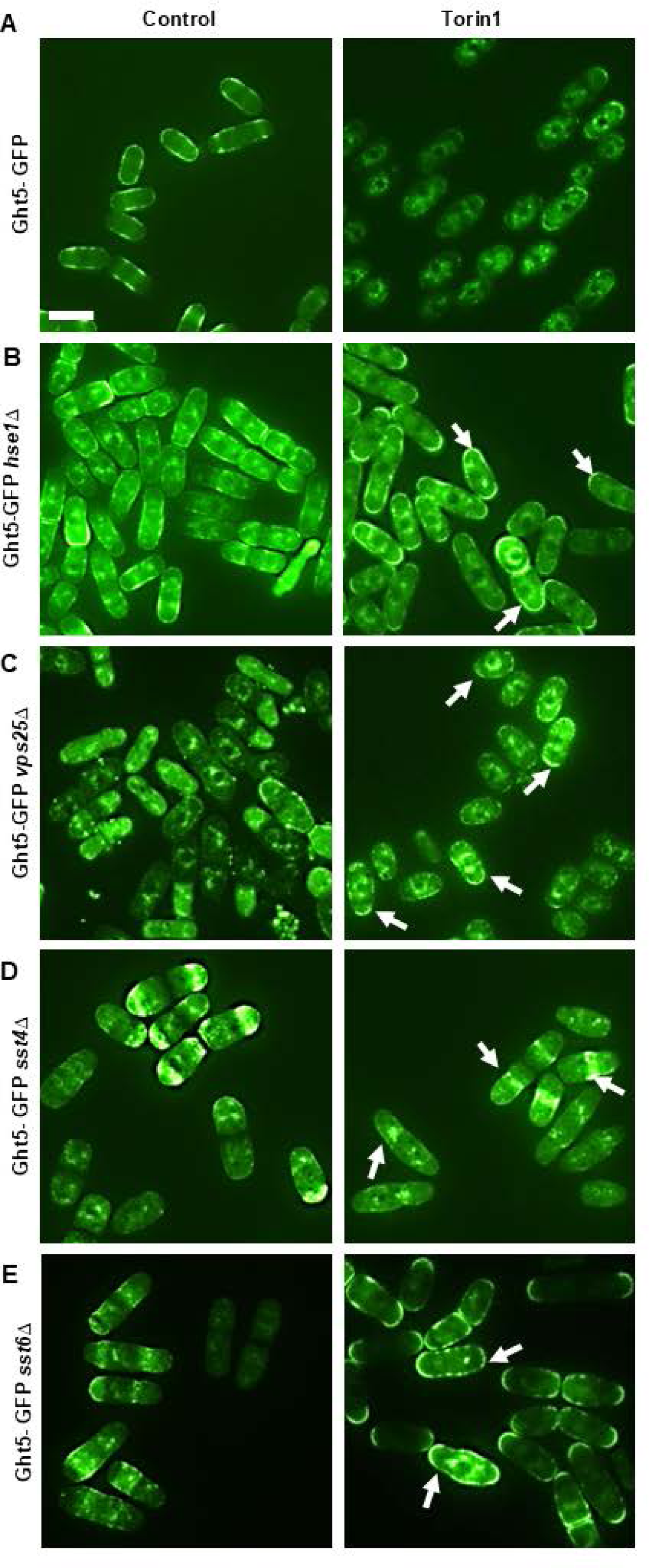
ESCRT mutants maintain Ght5 expression at the PM following exposure to Torin1. **A** *Wt* **B** *hse1*Δ, **C** *vps25*Δ, **D** *sst4*Δ and **E** *sst6*Δ strains were examined by fluorescent microscopy. Cells were exposed to 5µM Torin1 for 3h. White arrowheads indicate Ght5 expression at the PM. Scale bars indicate 10µm.

### Deletion of *pub1* abolishes Torin1- induced Aat1 degradation

TORC1-mediated ubiquitylation of the amino acid transporter Aat1 by Pub1 is required for its retention at the Golgi/ endosomes (ref). We investigated the effect of Torin1 on Aat1 in wt and *pub1*Δ mutants. As observed with Ght5-GFP, exposure to 5 µM Torin1 and nitrogen withdrawal induced the degradation of Aat1-GFP. The effect was not observed following exposure to caffeine and rapamycin, which induced Aat1-GFP degradation (Fig. 4A). This effect of Torin1 on Aat1 was abolished when *pub1* was deleted (Fig. 4B and 4D). Unlike Ght5-GFP, exposure to caffeine and rapamycin did not induce Aat1-GFP degradation (Fig. 1C and 4A). Interestingly, exposure to rapamycin induced the accumulation of Aat1-GFP in *pub1*Δ mutants. TORC1 negatively regulates Aat1 via Isp7 at the transcriptional level (12). Inhibition of TORC1 with rapamycin may induce *aat1* expression and accumulation of Aat1 in *pub1* mutants, as it cannot be degraded. (Fig. 4B). These findings also highlight significant differences between caffeine and rapamycin in terms of TORC1 inhibition (19,24). The deletion of *gsk3* did not abolish the Torin1-induced degradation of Aat1-GFP. Our observations suggest, however, that it does affect the dynamics of Aat1-GFP degradation following exposure to Torin1 (Fig. 4E and 4F).

**Fig. 4.**
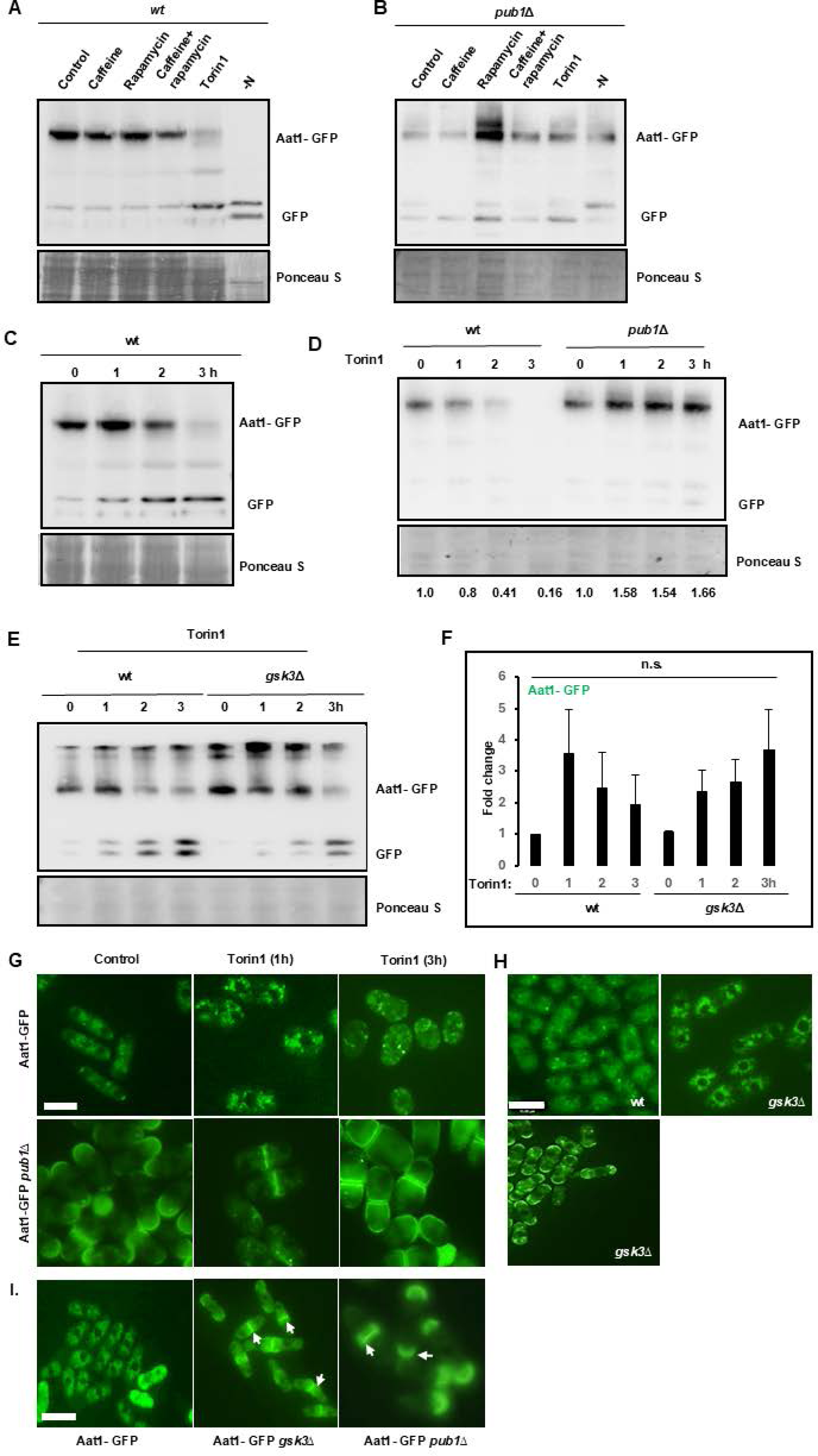
Torin1 induces Aat1 degradation in an ESCRT-dependent manner. **A** Cells expressing Aat1-GFP were exposed to 10mM caffeine, 100ng/mL Rapamycin alone or in combination, 5µM Torin1 or EMM2-N for 3h. Samples were analysed by immunoblotting. Full-length Aat1-GFP and cleaved GFP are indicated. **B** *pub1*Δ mutants expressing Aat1-GFP were treated as in A. **C** Wt cells expressing Aat1-GFP were exposed to 5µM Torin1 for the indicated timepoints and treated as in A. **D** *Wt* and *pub1*Δ mutants expressing Aat1-GFP were treated as in C. **E** *Wt* and *gsk3*Δ mutants expressing Aat1-GFP were treated as in C. **F** Quantification of Aat1-GFP expression in *wt* and *gsk3*Δ mutants exposed to 5µM Torin1 for 3h (n=3). **G** *Wt* and *pub1*Δ mutants expressing Aat1-GFP were exposed to 5µM Torin1 for 1 to 3h and examined by fluorescent microscopy. Scale bar indicates 10µm. **H** *Wt* and *gsk3*Δ mutants expressing Aat1-GFP were examined by fluorescent microscopy. Scale bar indicates 10µm. **I** *Wt, gsk3*Δ and *pub1*Δ mutants expressing Aat1-GFP were examined by fluorescent microscopy. White arrowheads indicate septa. Scale bar indicates 10µm.

We next examined the effect of Torin1 on Aat1 expression by fluorescence microscopy. In *wt* cells, exposure to Torin1 induced the transport of Aat1-GFP at the Golgi/ endosomes and subsequent degradation (Fig. 4G). Deletion of *pub1* resulted in the constitutive expression of Aat1-GFP at the PM as previously reported. Aat1-GFP also accumulated at the septa of dividing cells following exposure to Torin1. Pub1 has been reported to be partly localized to the septum in *S. pombe* (25). These findings demonstrate that Pub1 is required for the Torin1-induced degradation of Aat1-GFP (Fig. 4G). It was also observed that *pub1* mutants continued cell division, albeit asynchronously, following exposure to Torin1 (Fig. 4G). The larger cell size of *pub1*Δ mutants has been previously reported (25,26). Studies using Aat1-GFP *gsk3*Δ mutants produced variable results. In our experiments, the localization of Aat1-GFP in *gsk3*Δ mutants mimicked that observed in *pub1*Δ mutants (Fig. 4H). We noted, however, that Aat1-GFP exhibits a differential localization pattern in *gsk3*Δ mutants (Fig. 4G, 4H and I). These findings are consistent with previous reports linking TORC2 and Gsk3 to Pub1 stability and regulation of Aat1 expression (13). The inhibition of TORC2 induces Aat1 degradation. Furthermore, Pub1 but not Gsk3 is required for the Torin1-induced degradation of Aat1 in fission yeast. Both Pub1 and Gsk3 regulate Aat1 expression.

### Deletion of *pub1* abolishes Torin1- induced Cat1 degradation

As the PM amino acid transporter Cat1 is regulated like Aat1, we next investigated the effect of Torin1 on its expression. Exposure to 5 µM Torin1 induced Cat1-GFP degradation as observed in Aat1-GFP and Ght5-GFP expressing strains (Fig. 1, 2 and 5A). Cat1-GFP appeared to be more sensitive to exposure to 10 mM caffeine alone or with 100 ng/ mL rapamycin than Aat1-GFP (Fig. 5B). Deletion of *pub1*Δ abolished the effect of Torin1 on Cat1-GFP expression. We noted, however, that GFP still accumulated, indicating Pub1-independent alternate Cat1-GFP degradation mechanisms. Pub3 may thus substitute for Pub1 in mediating Cat1 but not Aat1 degradation (Fig. 5C). As observed with Aat1-GFP, deletion of *gsk3*Δ delayed but did not abolish Torin1-induced Cat1-GFP degradation in this strain (Fig. 5C and 5D). This observation resembles that observed with the *aat1-GFP* strain (Fig. 4E and 4F). Fluorescence microscopy revealed that exposure to Torin1 induced the relocation of Cat1-GFP from the Golgi/ endosomes to vacuoles. In untreated wt cells, Cat1 was localized to the Golgi/ endosomes. Exposure to Torin1 resulted in a diffuse appearance in the cytoplasm with bright speckles (Fig. 5Ea and 5Eb). Deletion of *pub1* resulted in the dispersion of Cat1-GFP from the Golgi/ endosomes to vacuoles to the cytoplasm, PM and septum of dividing cells (Fig. 5Ec and Ed). Deletion of *pub1Δ* prevented the degradation of Cat1-GFP following exposure to Torin1. While exposure to Torin1 in wt cells resulted in a speckled appearance, this was not observed in *pub1Δ* mutants, which fail to degrade Cat1 (Fig. 5B). Furthermore, *pub1* mutants continued to express Cat1-GFP at the PM following exposure to Torin1 (Fig. 5Ec and 5Ed). We next examined the effect of deleting *gsk3* in the *aat1-GFP, cat1-GFP, and Ght5-GFP* strains. The deletion of *gsk3* did not affect the localization of *cat1* -GFP as observed in pub1Δ mutants (Figure 5E and 5F). The deletion of *gsk3*, similarly, did not affect the expression of Ght5-GFP in untreated cells. Together, our findings suggest that while Pub1 regulates the PM expression of both Aat1 and Cat1, Gsk3 regulates only the localization of the former (Fig. 5F). This observation is consistent with previous reports linking Gsk3 to Pub1 activity and Aat1 expression. Pub1 is, however, required for the degradation of Aat1, Cat1 and Ght5 following exposure to Torin1. In this model, the deletion of *gsk3* would limit the availability of Pub1 and suppress sensitivity to Torin1 by maintaining nutrient transporter expression (13). Fission yeast cells can thus selectively fine-tune the expression of nutrient transporters under various conditions in a TORC1 and TORC2-dependent manner.

**Fig 5.**
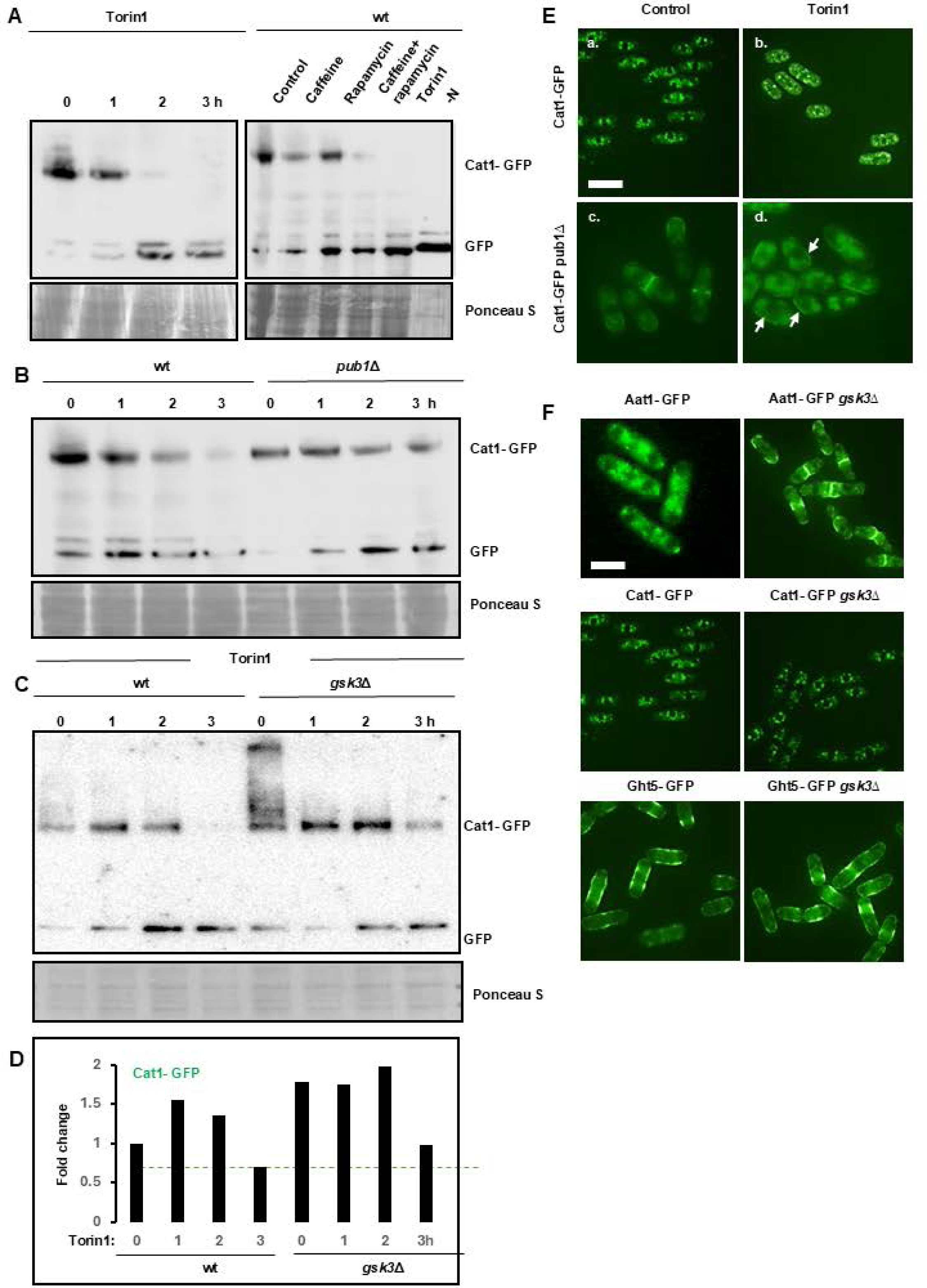
Torin1 induces Cat1 degradation in a Pub1-dependent manner. **A** Cells expressing Cat1-GFP were exposed to 5µM Torin1 for the indicated time points. Alternatively, the cells were exposed to 10mM caffeine, 100ng/mL Rapamycin alone or in combination, 5µM Torin1 or EMM2-N for 3h. Samples were analysed by immunoblotting. Full-length Cat1-GFP and cleaved GFP are indicated. **B** *Wt* and *pub1*Δ mutants expressing Cat1-GFP were exposed to 5µM Torin1 for the indicated timepoints. Samples were treated as in A. **C** *Wt* and *gsk3*Δ mutants expressing Cat1-GFP were exposed to 5µM Torin1 for the indicated timepoints. Samples were treated as in A. **D** Quantification of Aat1-GFP expression in *wt* and *gsk3*Δ mutants exposed to 5µM Torin1 for the indicated timepoints (n=3). **E** *Wt* and *pub1*Δ mutants expressing Cat1-GFP were exposed to 5µM Torin1 for 3h and examined by fluorescent microscopy. White arrowheads indicate expression of Cat1 at the PM. Scale bar indicates 10µm.

### The ubiquitin system is required for Torin1 sensitivity and links TORC1 and TORC2 signalling with cell division

Torin1 induces the ubiquitin-dependent degradation of nutrient transporters and Cdc25 by Pub1 (Fig. 1, 2, 3, 4 and 5) (19). E3 ligases such as Pub1 regulate multiple targets and pathways. Loss of Pub1 expression would prevent the degradation of Cdc25 and numerous nutrient transporters (9,9,10,19). The deletion of *pub1* did not confer resistance to caffeine combined with rapamycin or Torin1 but rather increased sensitivity to these drugs (Fig. 6A). Mutants lacking Pub1 are sensitive to various stresses and exhibit cell cycle defects, which may override its effect on Torin1 sensitivity (27). Gsk3 activity is elevated in *gad8*Δ mutants. These cells displayed increased sensitivity to caffeine combined with rapamycin or Torin1, consistent with TORC2 regulating sensitivity to these agents. As previously reported, *gsk3*Δ mutants are highly resistant to Torin1 (3). The deletion of *pub2* did not affect sensitivity to either caffeine plus rapamycin or Torin1. When both *pub2* and *pub3* were deleted, the cells exhibited resistance to Torin1 but not caffeine combined with rapamycin. A WD-repeat protein Lub1, the loss of which was found to show resistance to Torin1, is required for adequate ubiquitin expression in fission yeast, and *lub1*Δ mutants exhibit low global ubiquitylated protein levels (28). These mutants are highly resistant to Torin1, presumably due to their inability to sufficiently ubiquitylate target proteins (Fig. 6A). These findings identify a novel ubiquitin-dependent, ESCRT-regulated degradation system required for sensitivity to Torin1. We deleted *gsk3* in a strain expressing a Cdc25-GFP variant that lacks negative phosphorylation sites (29). In the Cdc25(12A)-GFP *gsk3*Δ mutant, expression of the phosphatase was extremely low and cleaved GFP was not detectable. Furthermore, exposure to Torin induced Cdc25(12A) degradation in wt but not *gsk3*Δ cells. Surprisingly, Torin1 induced Cdc25 accumulation in the absence of Gsk3 (Fig. 6B). This ESCRT-dependent system thus integrates nutrient transport with cell division through dual regulation of PM nutrient transporter and Cdc25 stability dependent on Pub1 (Fig. 6C).

**Fig 6.**
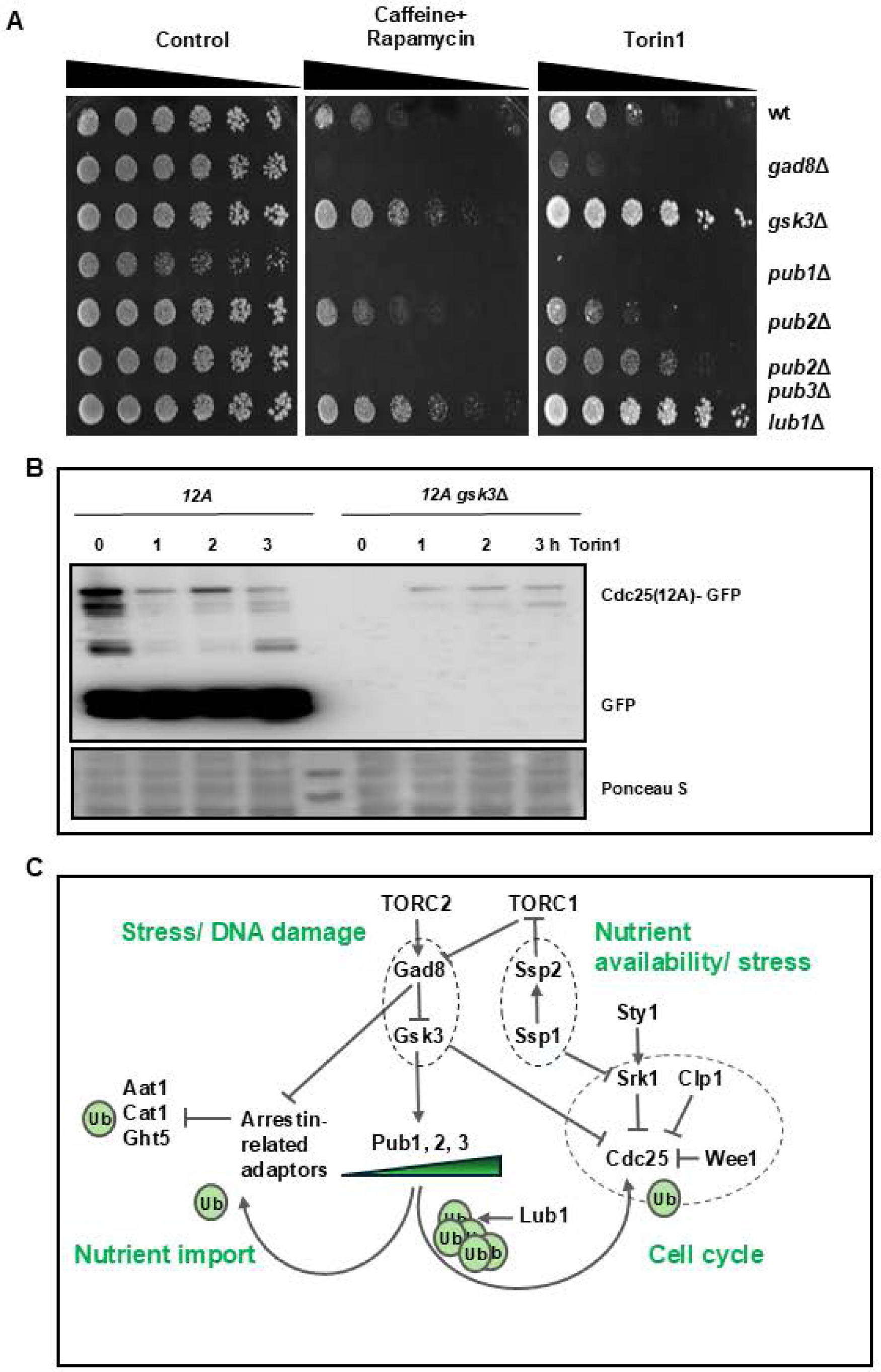
TORC2 and Pub1 integrate nutrient availability with cell division. **A** *Wt*, *gad8*Δ, *gsk3*Δ, *pub1*Δ, *pub2*Δ, *pub2*Δ *pub3*Δ strains were grown to stationary phase, diluted into fresh media at an O.D.600 of 0.3 and incubated for 2h. Cells were serially diluted and plated on YES media with 10mM caffeine and 100ng/mL rapamycin or 5µM Torin1 and incubated for 3-5 days. **B** *Wt* and *gsk3*Δ *Cdc25(12A)-GFP* strains were incubated with 5µM Torin1 for the indicated time points. Cell lysates were analyzed by immunoblotting. Full-length Cdc25(12A)-GFP and cleaved GFP are indicated. **C** TORC1 and TORC2 adjust nutrient transporter expression to cell cycle regulation. Environmental and nutrient stress inhibit TORC1-mediated Gad8 inhibition, resulting in Gsk3 activation. Gsk3 stabilises Pub1 which is required for the ubiquitylation and degradation of PM nutrient transporters. Gad8 inhibits the arrestin-related-mediated ubiquitylation of Ght5 by Pub1. Low endogenous ubiquitin levels prevent optimal target protein degradation. TORC2 also positively regulates Cdc25 stability and is required for cell cycle re-entry following activation of DNA damage checkpoints. TORC1 and TORC2 also interact with Ssp1 and Ssp2 which also regulate cell cycle kinetics.

## Discussion

We have identified an ESCRT-dependent protein degradation system that selectively regulates sensitivity to Torin1 in fission yeast. E3 ligases target nutrient transporters for degradation in lysosomes via the ESCRT pathway, which differs mechanistically from autophagy (4,30). TOR inhibitors have drawn considerable attention as anticancer agents, due to their central role in cellular metabolism, autophagy and cell division (1,2). Their use as anticancer agents has been limited by modest anti-tumor activity and various drug resistance mechanisms. Current studies investigate the impact of cell metabolism on the inhibitory activities of these drugs and the mechanisms that underlie resistance to them (2). We present evidence that suppression of Torin1-induced degradation of nutrient transporters is sufficient for resistance to this drug. To our knowledge, such a resistance mechanism has not been previously reported for TOR inhibitors. Clinically, deregulated ESCRT signalling has been proposed as a potential mechanism of drug resistance. Compromised EGFR sorting to lysosomes via the ESCRT pathway could potentially result in resistance to EGF receptor signalling. Thus, drug resistance need not be necessarily linked to mutations altering the drug affinity of a target protein or the activity of downstream effectors. Our findings in an evolutionarily conserved model system confirm the need for further investigations into deregulated ESCRT signalling as a mechanism of TORC inhibitor drug resistance (6,31). Crucially, this system does not mediate Torin1 resistance due to a single deregulated protein. Rather, compromised individual proteins at any stage of the ESCRT-mediated degradation pathway, including low ubiquitin levels, confer resistance to Torin1. We identified only *atg6* and *vps38* mutants as exhibiting Torin1 resistance in the study by Rodríguez-López *et al*., (27). Atg6 and Vps38 direct protein degradation via the ESCRT pathway, while Atg38 regulates autophagy. We also noted that the majority of *atg* gene deletions strongly enhanced sensitivity to Torin1 (21). Thus, defective ESCRT signalling but not autophagy induces resistance to Torin1 (Fig. 1A).

TORC1 and 2 regulate the expression of nutrient transporters through the ESCRT system, which is dependent on the E3 ligases Pub1 and Pub3 (10,11,13,32). Cells ensure the adequate import of nutrients by selectively adjusting the expression levels of nutrient transporters, including Aat1, Cat1 and Ght5. On nitrogen withdrawal, membrane transporters relocate to the PM and/ or are directly degraded via the ESCRT system. Glucose deprivation induces G2 arrest, and both conditions lead to cell cycle exit and induction of autophagy (11,32–34). TORC2 positively regulates protein expression levels by preventing their degradation by the ESCRT system. Exposure to Torin1 but not caffeine or rapamycin induces cell cycle arrest. Thus, Torin1 inhibits fission yeast proliferation by inhibiting TORC2. We have demonstrated that exposure to Torin1 mimics nitrogen withdrawal by inducing the degradation of amino acid and glucose transporters. This degradation requires the prior arrestin-related adaptor-dependent ubiquitylation of substrate proteins by Pub1 and Pub3 (10,11,25). We have shown that mutants defective in ESCRT-mediated degradation display strong resistance to and fail to degrade Ght5 in response to Torin1. Hence, Torin1 sensitivity correlates strongly with uncompromised ESCRT signalling. ESCRT mutants such as *hse1*Δ and *vps25*Δ fail to degrade Ght5 following exposure to Torin1 and are highly resistant to the drug. Aat1, Cat1 and Ght5 mutants lacking *pub1* fail to degrade these transporters following Torin1 exposure. Indeed, the degradation of Aat1 does not occur in response to nitrogen withdrawal in *pub1*Δ mutants. In contrast to *wt* cells, therefore, ESCRT and *pub1*Δ mutants continue to express nutrient transporters at the PM. This may permit the sufficient importation of nutrients to prevent cell cycle exit. The deletion of *pub*1 did not confer resistance to Torin1. This may reflect its dual role in negatively regulating Cdc25. We noted that Torin1 failed to arrest but induced asynchronous cell division in these mutants. Interestingly, low endogenous ubiquitin levels are also associated with Torin1 resistance. In these mutants, insufficient ubiquitin availability prevents the degradation of target proteins. Mutants that fail to de-ubiquitylate target proteins are also resistant to this agent, likely due to ubiquitin sequestration by target proteins, which accumulate within the cell or on the PM (3,7). Our study identifies an ESCRT system that includes proteins maintaining ubiquitin homeostasis, arrestin-related adaptors, Pub1 and Pub3, and vacuolar biosynthesis whose expression is required for Torin1 sensitivity. TORC1 and 2 integrate the availability of nutrients with cell division through this system by co-regulation of nutrient transporters and Cdc25 (ref, ref) (Fig. 6C)

While caffeine in combination with rapamycin does slow cell division, our studies suggest that the underlying mechanism differs substantially from that of Torin1. Torin1 directs Ght5 for degradation through the ESCRT system. In contrast, caffeine plus rapamycin induces the degradation of Cat1 and Ght5 but not Aat1. Significantly, rapamycin induced Aat1 accumulation in *pub1*Δ mutants. We noted that cleavage of Aat1-, Cat1- and Ght5-GFP still occurs in *pub1* mutants. Furthermore, ESCRT mutants activate autophagy in the presence of Torin1. Caffeine combined with rapamycin may target Cat1 and Ght5 for degradation via autophagy. The co-deletion of *pub2* and *pub3* conferred resistance to Torin1 but not caffeine and rapamycin. Low ubiquitin expression in *lub1* mutants presumably induces Torin1 resistance by suppressing Pub1 and Pub3 activity (28). These mutants exhibited resistance to caffeine plus rapamycin and Torin1, but there is no correlation in drug resistance between them. The ESCRT system thus mediates sensitivity to Torin1 but not caffeine when combined with rapamycin. Caffeine has been shown to indirectly inhibit TORC1 by inducing the activation of AMPK. These findings suggest that caffeine, a well-characterised drug, can suppress resistance to TOR inhibitors. Caffeine in combination with rapamycin may trigger nutrient transporter degradation independently of ESCRT signalling. We previously reported that mutants unable to transfer substrate proteins from the Golgi/ endosomes to lysosomes are also resistant to Torin1. Future studies will investigate the role of the ESCRT system in mediating sensitivity to Rapalink-1 (3,35).

## Conclusions

We have identified an ESCRT-dependent pathway required for sensitivity to Torin1. A role for this pathway in mediating resistance to TOR inhibitors has not been previously reported. Mutants unable to degrade PM nutrient transporters continue to undergo cell division in the presence of Torin1. Our findings partly explain why Torin1, but not caffeine, combined with Rapamycin induces cell cycle arrest. Crucially, ubiquitin homeostasis is similarly required for sensitivity to Torin1.

## List of abbreviations

EGFR: Epidermal Growth Factor Receptor
ESCRT: Endosomal Sorting Complex Required for Transport
GFP: Green Fluorescent Protein
PM: Plasma Membrane
TORC1: Target of Rapamycin Complex 1
TORC2: Target of Rapamycin Complex 2

## Declarations

### Consent for publication

All authors consented to the publication of the manuscript.

## Availability of data and materials

Data and yeast strains are available on request, subject to permission of the creators.

## Competing interests

The authors declare no competing interests.

## Funding

C.R. acknowledges funding from the Royal Society (Research Award grant number RGS∖R1∖201348), BBSRC (grant numbers BB/V006916/1, BB/V006916/2) and MRC (grant number MR/W001462/1). Y.T. acknowledges funding from Grants-in-Aid for Scientific Research (C) from the Japan Society for the Promotion of Science (23K05758).

## Authors’ contributions

J.P.A. and C.R. designed the study. J.P.A. and R.A.I. carried out experiments. Y.T. prepared the yeast mutants. J.P.A., Y.T. and S.S. and C.R. analyzed the data. J.P.A. wrote the manuscript. J.P.A., R.A.I., C.R., Y.T. and S.S. revised and edited the manuscript

## Acknowledgements

We are grateful to the Matsumoto and Nakashima laboratories and the Yeast Genetic Resource Centre (YGRC) Japan for mutant *S. pombe* strains.

---

**Additional file 1.**
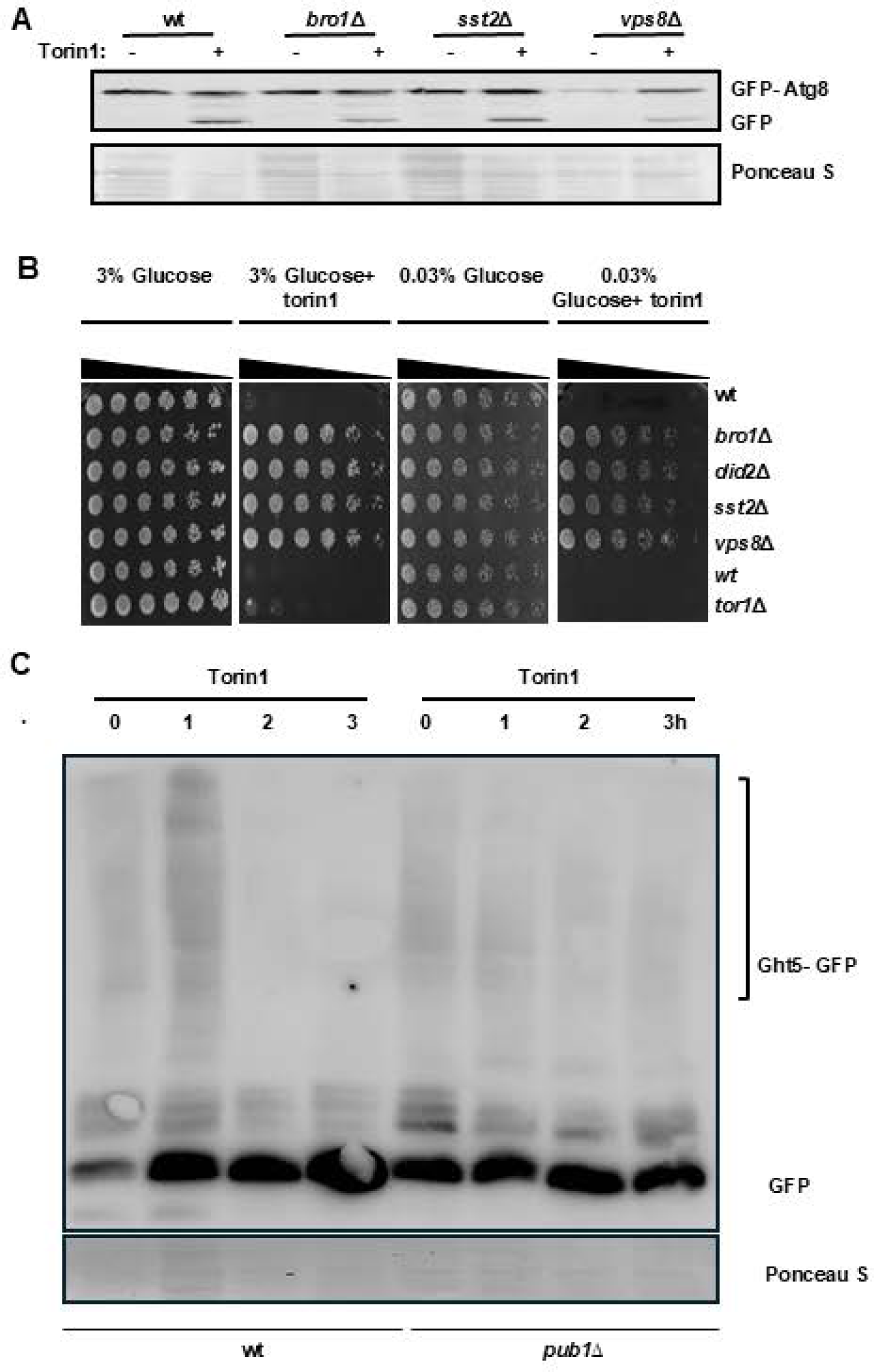
A Wt, *bro1*Δ, *sst2*Δ and *vps8*Δ strains expressing GFP-Atg8 were grown to log phase and exposed to 5µM Torin1 for 2h. Cell pellets were resolved by SDS-PAGE and blotted with monoclonal antibodies directed against GFP. **B** Wt, *bro1*Δ, *did2*Δ, *sst2*Δ and *vps8*Δ strains were grown to stationary phase, diluted into fresh media and incubated for a further 2h. Cells were serially diluted and plated on YES media with 3% or 0.03% glucose with or without 5µM Torin1 and incubated for 3-5 days.

**Additional Table 1.** *S. pombe* strains used in this study.

| Strain | Genotype | Source/Reference |
| --- | --- | --- |
| *h<sup>-</sup>* L972 | *h<sup>-</sup>* | Laboratory stocks |
| *tor1Δ* | *h<sup>-</sup> tor1::kanMX6* | Laboratory stocks/(Rallis *et al.*, 2013) |
| *gad8Δ* | *h<sup>-</sup> gad8::ura4 ade6- M216 leu1 ura4- D18* | YGRC |
| *gsk3Δ* | *h<sup>-</sup> gsk3::kanMX6* | Laboratory stocks/ (Rallis *et al.*, 2017) |
| *lub1Δ* | *h<sup>+</sup> lub1::KanMX ade6-M216 ura4-D18 leu1-32* | Laboratory stocks |
| *ght5-GFP* | *h<sup>-</sup> gad8<sup>TS</sup> ght5-GFP::KanR* | Laboratory stocks/ (Saitoh *et al.*, 2015) |
| *ght5-GFP gad8<sup>TS</sup> gsk3Δ* | *h<sup>-</sup> gad8<sup>TS</sup> gsk3::natMX ght5-GFP::KanR* | Laboratory stocks/ (Toyoda *et al.*, 2021) |
| *ght5-GFP gad8<sup>TS</sup> pub1Δ* | *h<sup>-</sup> gad8<sup>TS</sup> pub1::hphR ght5-GFP::KanR* | Laboratory stocks/ (Toyoda *et al.*, 2025) |
| *ght5-GFP gad8<sup>TS</sup> hse1Δ* | *h<sup>-</sup> gad8<sup>TS</sup> hse1::hphR ght5-GFP::KanR* | Laboratory stocks/ (Toyoda *et al.*, 2021) |
| *ght5-GFP gad8<sup>TS</sup> vps25Δ* | *h<sup>-</sup> gad8<sup>TS</sup> vps25::hphR ght5-GFP::KanR* | Laboratory stocks/ (Toyoda *et al.*, 2021) |
| *ght5-GFP gad8<sup>TS</sup> vps28Δ* | *h<sup>-</sup> gad8<sup>TS</sup> vps28::hphR ght5-GFP::KanR* | Laboratory stocks/ (Toyoda *et al.*, 2021) |
| *ght5-GFP gad8<sup>TS</sup> sst4Δ* | *h<sup>-</sup> gad8<sup>TS</sup> sst4::hphR ght5-GFP::KanR* | Laboratory stocks/ (Toyoda *et al.*, 2021) |
| *ght5-GFP gad8<sup>TS</sup> sst6Δ* | *h<sup>-</sup> gad8<sup>TS</sup> sst6::hphR ght5-GFP::KanR* | Laboratory stocks/ (Toyoda *et al.*, 2021) |
| *aat1-GFP* | *h<sup>-</sup> aat1-GFP::KanR* | Matsumoto Laboratory |
| *aat1-GFP gsk3Δ* | *h<sup>-</sup> aat1-GFP<LEU2 leu1-32 gsk3::KanMx6* | This study |
| *aat1-GFP pub1Δ* | *h<sup>-</sup> pub1::ura4+ ura4-D18 aat1-GFP<<LEU2 leu1-32* | Matsumoto Laboratory |
| *cat1-GFP* | *h<sup>-</sup> cat1+-GFP::kanR* | Nakashima laboratory |
| *cat1-GFP gsk3Δ* | *h- gsk3::natMX cat1+-GFP-kanMX* | This study |
| *cat1-GFP pub1Δ* | *h- pub1::hphMX cat1+-GFP-kanMX* | Nakashima laboratory |
| *cdc25 (12A)- GFP* | *h- cdc25(12A)-GFPint cdc25:: kanMX6 ura4-D18 leu1-32* | Young laboratory |
| *cdc25 (12A)- GFP*<br>*gsk3Δ* | *h- cdc25(12A)-GFPint cdc25:: kanMX6 gsk3::natMX6 ura4-D18 leu1-32* | This study |
| *pub1Δ* | *h- pub1::ura4+ leu1-32 ura4-D18* | YGRC |
| *pub2Δ* | *h+ pub2::ura4+ ura4-D18* | YGRC |
| *pub2Δ pub3Δ* | *h90 pub2::ura4+ pub3::ura4+ ade6-M216 leu1-32 ura4-D18* | YGRC |
| *GFP-atg8* |  | Laboratory stocks |
| *GFP-atg8 bro1Δ* |  | Laboratory stocks |
| *GFP-atg8 sst2Δ* |  | Laboratory stocks |
| *GFP-atg8 vps8Δ* |  | Laboratory stocks |

\*Yeast Genetic Resource Center (YGRC) Japan

## Notes

### Competing Interest Statement

The authors have declared no competing interest.

### Summary of Updates

Figure 1 is revised. The revision is in panel 1E in order to remove a duplicated western blot. The updated figure now contains the right western blot. This update does not change the results and conclusions of the study.

